# The evolutionary dynamics of the odorant receptor gene family in corbiculate bees

**DOI:** 10.1101/130781

**Authors:** Philipp Brand, Santiago R. Ramírez

## Abstract

Insects rely on chemical information to locate food, choose mates, and detect potential predators. It has been hypothesized that adaptive changes in the olfactory system facilitated the diversification of numerous insect lineages. For instance, evolutionary changes of Odorant Receptor (OR) genes often occur in parallel with modifications in life history strategies. Corbiculate bees display a diverse array of behaviors that are controlled through olfaction, including varying degrees of social organization, and manifold associations with floral resources. Here we investigated the molecular mechanisms driving the evolution of the OR gene family in corbiculate bees in comparison to other chemosensory gene families. Our results indicate that the genomic organization of the OR gene family has remained highly conserved for approximately 80 million years, despite exhibiting major changes in repertoire size among bee lineages. Moreover, the evolution of OR genes appears to be driven mostly by lineage-specific gene duplications in few genomic regions that harbor large numbers of OR genes. A selection analysis revealed that OR genes evolve under positive selection, with the strongest signals detected in recently duplicated copies. Our results indicate that chromosomal translocations had a minimal impact on OR evolution, and instead local molecular mechanisms appear to be main drivers of OR repertoire size. Our results provide empirical support to the longstanding hypothesis that positive selection shaped the diversification of the OR gene family. Together, our results shed new light on the molecular mechanisms underlying the evolution of olfaction in insects.

## Introduction

Animals have evolved sophisticated sensory systems that can detect and discriminate airborne volatile chemicals and provide precise information about food, enemies, and mating partners (Hansson & Stensmyr 2011). Insects detect olfactory signals and cues via olfactory sensory neurons located on the antenna. Each olfactory sensory neuron expresses chemosensory receptor proteins on the cell membrane (Vosshall et al. 2000; Elmore & D P Smith 2001; Dobritsa et al. 2003). It is the interaction of volatile odorant molecules with these receptor proteins that initiates the signal transduction and transmission toward the olfactory centers in the insect brain (Vosshall et al. 2000; Dobritsa et al. 2003). Insect genomes are endowed with a diverse array of functional olfactory chemosensory receptor genes, each of which encodes a unique receptor protein tuned to a specific set of odors (Hallem & Carlson 2006; Hallem et al. 2004; Wang et al. 2010; Benton et al. 2009). The entire repertoire of receptor genes expressed in the olfactory organs determines the spectrum of chemical volatiles that an insect species can detect. The majority of olfactory chemosensory receptors in insects belong to the Odorant Receptor (OR) gene family, but also include Ionotropic Receptors (IRs) (Sánchez-Gracia et al. 2009; Croset et al. 2010).

Comparative analyses of insect genomes have revealed that the number of OR genes and divergence among them can vary widely between species. Insect genomes may contain as few as 10 OR genes (head lice, (Kirkness et al. 2010)) and as many as 300 OR genes (ants, (Christopher D Smith et al. 2011; Chris R Smith et al. 2011)). In fact, the expansion and contraction of the OR gene family has been linked to shifts in the sensory ability of some insect lineages. This supports the idea that adaptation to novel food resources (McBride 2007; McBride & Arguello 2007; Goldman-Huertas et al. 2015) or modifications in the pheromone communication system (Gould et al. 2010) can be mediated by peripheral changes in the OR gene repertoire. In addition, changes in the amino acid sequence of existing OR genes have been shown to correspond to adaptive shifts in sensory tuning (Leary et al. 2012; Pellegrino et al. 2011). However, our current understanding of the genetic and molecular mechanisms that drive diversification of the insect OR gene family is derived from a few insect lineages (Nei et al. 2008; Sánchez-Gracia et al. 2009). Thus, the inclusion of more lineages is needed to obtain a better picture of the general and lineage-specific patterns of OR gene family evolution.

Gene family evolution is determined by multiple molecular mechanisms including genomic drift, natural selection, and chromosomal rearrangements. Birth-death processes can directly impact the number of genes within a gene family (Nei & Rooney 2005). For example, genomic drift may generate new gene copies through gene duplication (gene ‘birth’) while existing copies can be purged via pseudogenization or deletion (gene ‘death’). Novel mutations resulting in gene duplication or gene loss may be subsequently fixed through neutral genetic drift or positive selection (Nei & Rooney 2005; Nei et al. 2008; Innan & Kondrashov 2010). Although OR gene repertoire sizes have been analyzed extensively in various insect lineages, the relative contribution of positive selection and neutral processes in a birth-death evolutionary framework remains uncertain (Nei et al. 2008; Sánchez-Gracia et al. 2009).

The genomic organization and architecture of gene families may contain signatures of past evolutionary forces that shaped their diversification. For instance birth-death processes often produce tandem arrays of OR genes, which are groups of ancestrally duplicated genes located in close physical proximity along the genome (Robertson & Wanner 2006; Nei & Rooney 2005). Chromosomal rearrangements and transposition, on the other hand, can lead to inter-chromosomal translocations of OR genes (Guo & Kim 2007; Conceição & Aguadé 2008), and it has been hypothesized that changes in the location of OR genes along the genome can influence their evolution (Sánchez-Gracia et al. 2009; Kratz et al. 2002; Conceição & Aguadé 2008). Although both tandem duplications and chromosomal rearrangements may influence the evolution of the OR gene family, the relative contribution of each process remains poorly understood in insects and surprisingly few studies have addressed this problem in the OR gene family (Sánchez-Gracia et al. 2009).

Like most insects, bees rely on olfactory information to regulate a wide array of behaviors, including the location of food sources and nesting materials, the identification of mating partners, and social interactions with other colony members in social species. Corbiculate bees encompass a group of ecologically and economically important bee lineages, including honey bees, bumble bees, stingless bees, and orchid bees. Honey bees are estimated to pollinate one-third of agricultural crops (Klein et al. 2007), and stingless bees and orchid bees are the major pollinators of numerous tropical flowering plant species (Slaa et al. 2006; Ramírez et al. 2002). Corbiculate bees have evolved a variety of phenotypic and behavioral traits that require specialized olfactory functions. These include the evolution of an obligate cooperative social lifestyle (eusociality), which has resulted in the evolution of specialized pheromone communication systems in honey bees, bumble bees and stingless bees (Grüter & Keller 2016; Kocher & Grozinger 2011; Leonhardt et al. 2016; Leonhardt 2017). In addition, orchid bees evolved a unique pheromone communication system in which male bees concoct species-specific perfume bouquets from scents collected from flowers and other sources to subsequently use during courtship display (Vogel 1966; Eltz et al. 1999; Roubik & Hanson 2004). However, despite the central role that olfactory encoding plays in the biology of corbiculate bees, the OR gene family has been studied only in a limited number of bee lineages (Robertson & Wanner 2006; Sadd et al. 2015; Brand et al. 2015; Park et al. 2015).

Here we analyzed the evolution of the OR gene family in comparison to other chemosensory gene families in a set of species including the four corbiculate bee clades, spanning 80 million years (my) of evolution. We tested whether the evolution of the OR gene family has been influenced by changes in the genomic organization of genes, as hypothesized based on patterns observed in *Drosophila* and mammalian genomes (Sánchez-Gracia et al. 2009; Kratz et al. 2002; Conceição & Aguadé 2008). Second, we tested the hypothesis that birth-death processes, in combination with positive selective pressures, promoted the origin of novel OR genes. Finally, we estimated the relative importance of positive selection in shaping OR sequence divergence, a mechanism that has been hypothesized to drive sensory adaptation in insect lineages (Brand et al. 2015; Leary et al. 2012; McBride et al. 2014).

## Results

### The Evolution of Corbiculate Bee Odorant Receptors is Highly Dynamic

To analyze the evolutionary dynamics of odorant receptor (OR) gene repertoires in corbiculate bees, we reconstructed the phylogenetic relationships among the entire OR gene family based on both whole-genome sequences and transcriptome data obtained from ten bee species (Fig. 1). An initial comparison of OR annotation based on genomic and transcriptomic data indicated no differences in annotation accuracy, but as expected, the number of genes tended to be lower for transcriptome data (Supplemental Text; (Grosse-Wilde et al. 2011; Brand et al. 2015)). Our analysis included published whole-genome sequence data from at least one species per corbiculate bee tribe (Robertson & Wanner 2006; Weinstock et al. 2006; Sadd et al. 2015; Kapheim et al. 2015) as well as the published antennal transcriptome of the orchid bee *Euglossa viridissima* (Brand et al. 2015). In addition, we sequenced the genome of one orchid bee (*Euglossa dilemma;* (Brand et al. 2017)) and the antennal transcriptomes of four additional orchid bee species. The resulting dataset, which included three orchid bee species pairs, allowed us to analyze the evolution of the OR gene family at multiple taxonomic levels over a broad range of divergence times (0.15 to 80mya; Fig. 1). We compared the evolution of the OR gene family, the largest and most dynamic chemosensory gene family in insects, to the evolution of four additional chemosensory gene families, including the Gustatory Receptors (GRs), Ionotropic Receptors (IRs), Odorant Binding Proteins (OBPs), and Chemosensory Proteins (CSPs). We used genomic data only to derive evolutionary patterns of gene family size since transcriptome data often yield incomplete gene sets (Fig. 1).

**Fig. 1.**
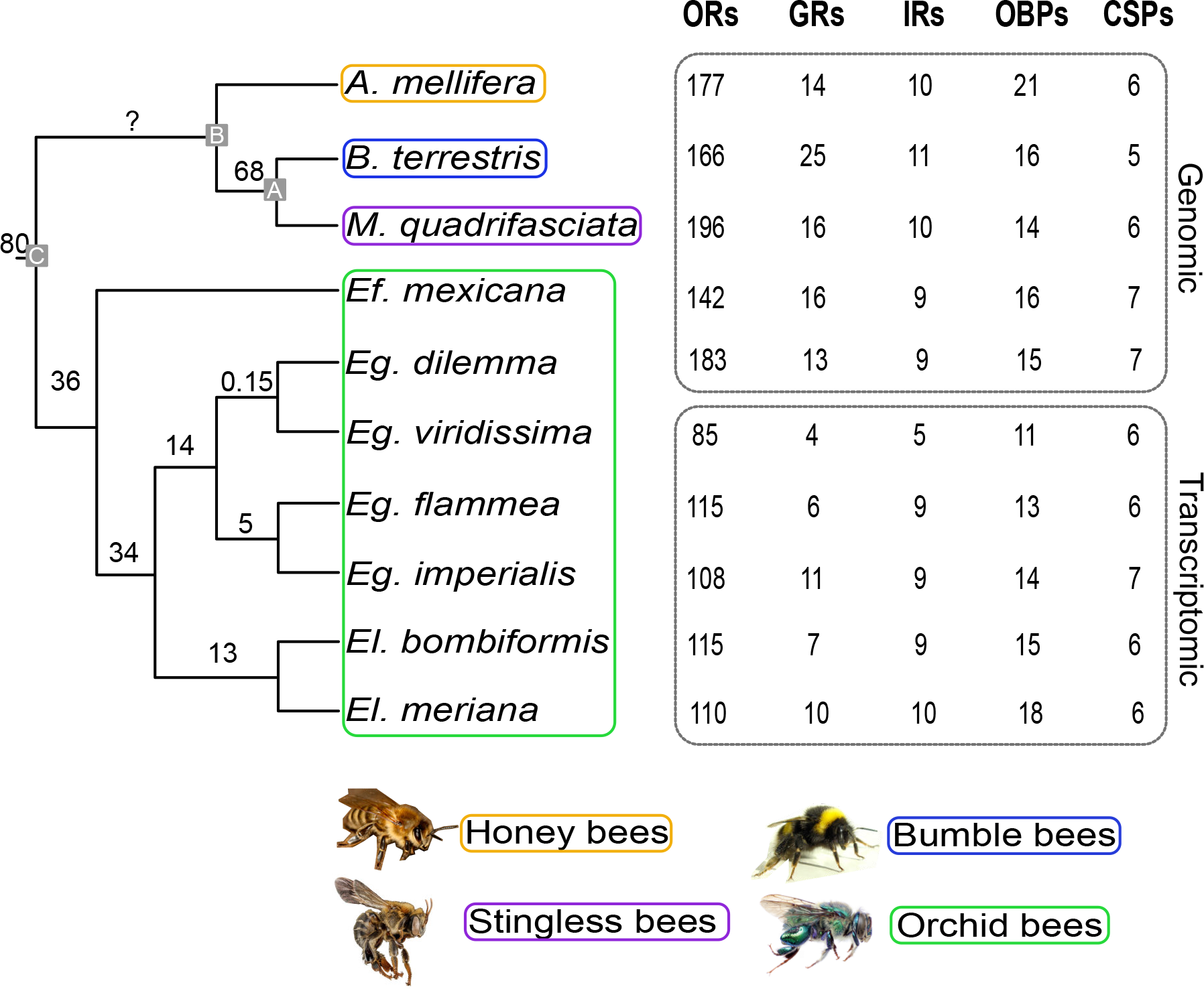
Gene family size dynamics in corbiculate bees. Numbers of previously (*A. mellifera (Robertson & Wanner 2006), B. terrestris (Sadd et al. 2015), Eg. viridissima (Brand et al. 2015)*) and newly annotated (all others) genes for each gene family are indicated. Annotations derived from whole genomes and antennal transcriptomes are labeled. Phylogenetic relationships and divergence times of the ten corbiculate bee species following Ramirez et al. 2010 and Romiguier et al. 2015 are shown. Annotated nodes indicate MRCA of bumble bees and stingless bees (A), honey bees + A (B) and all corbiculate bees (C).

Gene repertoire size varied substantially among bee lineages, ranging from 142 OR genes in the orchid bee *Eufriesea mexicana* to 196 OR genes in the stingless bee *Melipona quadrifasciata* (Fig. 1). The identified ORs were organized in 115 homologous OR ortho-groups consisting of orthologs and/or inparalogs (*i.e*. lineage-specific expansions, Fig. 2, Fig. S1). Of these ortho-groups, 24 did not contain duplications but instead consisted of simple 1:1 orthologous genes that were present in three or more bee species (see Fig. 2b for an example). The remaining 91 ortho-groups included species-specific, genus-specific, and tribe-specific duplications or larger expansions (see Fig. 2c for an example). In contrast to the other four gene families analyzed, the repertoire size of ORs is highly variable among corbiculate bee lineages. The majority of genes in the non-OR gene families revealed simple 1:1 orthology with few lineage-specific duplications resulting in similar repertoire sizes among bee lineages (Fig. 1; Fig. S2). An exception to this was an observed increase in gene family size in honey bee OBPs as well as bumble bee GRs, supporting previous observations (Supplemental Text; (Forêt & Maleszka 2006; Sadd et al. 2015). Among all chemosensory genes, we identified 32 gene duplications and large expansions in the honey bees, 36 in the bumble bees, 26 in the stingless bees, and 18 in the orchid bees, most of which were observed in the OR gene family (89%, Tab. S1). These results show that the evolution of the corbiculate bee OR gene family repertoire is characterized by highly dynamic lineage-specific gene expansions.

**Fig. 2.**
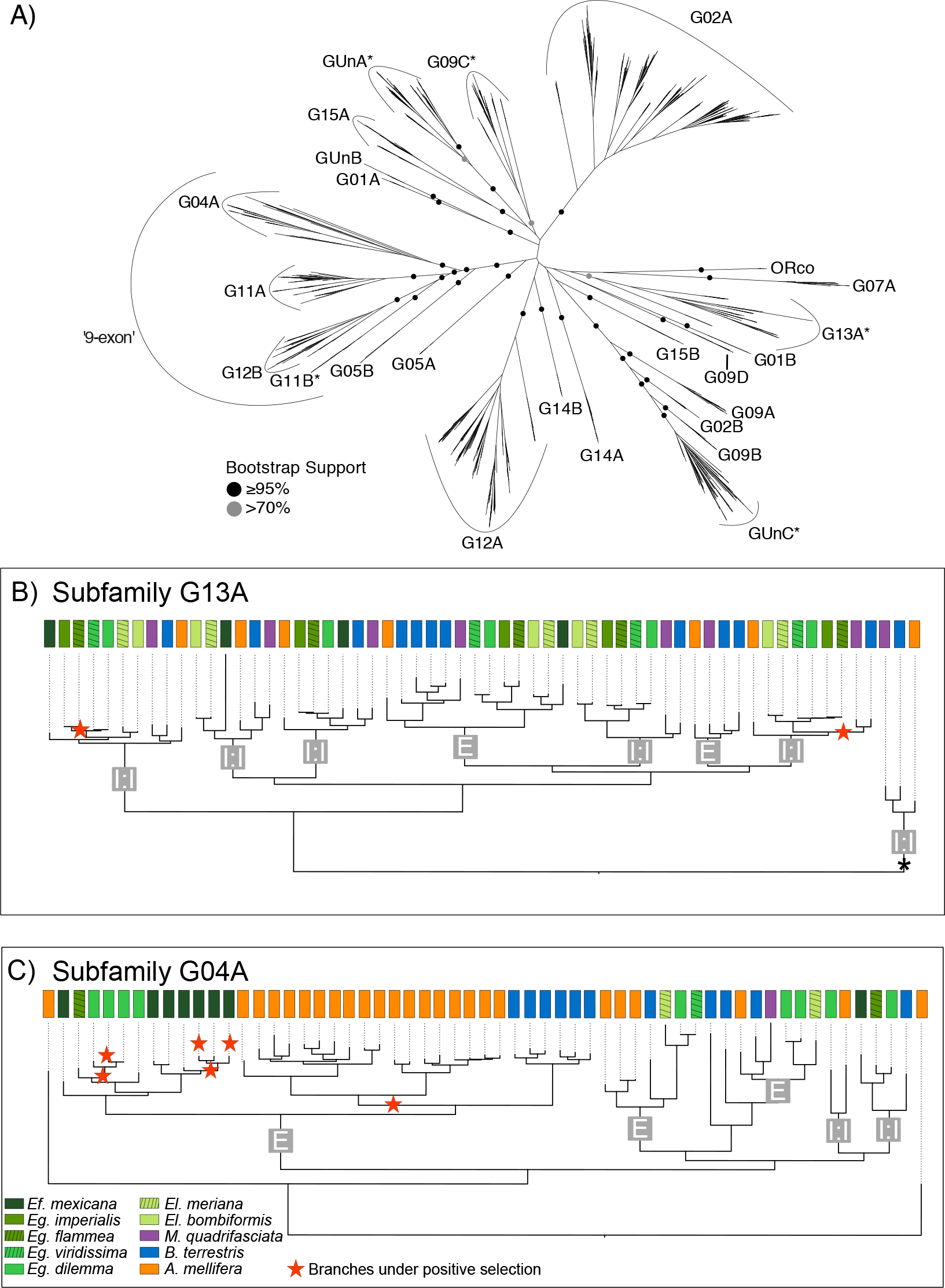
Odorant receptor subfamily dynamics in corbiculate bees. Relationships between subfamilies (**A**) are based on all annotated ORs of 10 corbiculate bee species. Observed differences of within subfamily dynamics are shown with subfamily G13A and subfamily G04A as examples (**B, C**). (**A**) OR subfamilies have high bootstrap support while deeper nodes are of low support. All genes within a subfamily are located in a single tandem array with highly conserved genomic landscapes throughout corbiculate bees. The few exceptions are indicated with *. (**B**, ORs in subfamily G13A are mainly simple 1:1 single copy orthologs found in all or a subset of the analyzed species (marked ‘1:1’) in contrast to subfamily G04A, where lineage specific duplications and larger expansions in *A. mellifera*, *B. terrestris*, *Eg. dilemma* and *Ef. mexicana* dominate (marked ‘E’). Branches evolving under positive selective pressures are indicated by a red star. Both, duplicate branches as well as branches in 1:1 ortholog groups were under positive selection. The branch indicated with a * in (**B**) indicates the sistergroup to all other ORs in subfamily G13A which is found at a genomic location different from the rest of the subfamily in all corbiculate bee species representing an ancient translocation event (see text for details).

### The Genomic Organization of the OR Gene Family is Highly Conserved in Corbiculate Bees

Next, we investigated the genomic organization of OR genes across the corbiculate bees. Our analysis revealed that OR genes are widely distributed throughout the five bee genomes we analyzed. This supports previous studies where OR genes were detected on scaffolds assigned to almost all chromosomes in those species with available linkage groups (*i.e*. honey bees and bumble bees; (Weinstock et al. 2006; Sadd et al. 2015; Robertson & Wanner 2006)). Congruent with a birth-death process for OR gene evolution, the majority of genes were clustered in large tandem arrays and only 16 ORs were found in isolated genomic regions as singletons. We tested whether the chromosomal location of OR genes is conserved among corbiculate bees by comparing the 200 kb flanking regions of all OR singletons and tandem arrays among species using an “all-against-all” reciprocal blastn approach.

With the exception of a small number of ancestral translocations (Supplementary Text), we found that the flanking regions and thus the physical chromosomal locations of most OR genes were strictly homologous in all corbiculate bee species, including 12 of the 16 singletons detected (Tab. S2). Furthermore, all orthologous OR genes found in two or more species were located in homologous chromosomal regions. Paralogous genes within species-specific, genus-specific, and tribe-specific duplications and larger OR sublineage expansions were always located in the same tandem array. Similarly, ORs located within the same genomic region were generally more closely related to each other than ORs located in different genomic regions that formed highly supported monophyletic gene clades (Fig. 2a). On the basis of this strong correlation between phylogenetic relationship and genomic location, we defined a total of 25 well-supported subfamilies in the OR gene family that largely corresponded to groups previously identified in ants (Fig. 2a; (Zhou et al. 2012; Engsontia et al. 2015)). In addition to the high conservation of genomic locations of OR genes and tandem arrays, we found that the organization of genes within tandem arrays (*i.e*. gene microsynteny) is also highly conserved (Supplementary Text; Fig. S3). Our analysis identified similar patterns of conservation in the genomic organization of other chemosensory gene families, including Gustatory Receptors, Ionotropic Receptors, Odorant Binding Proteins, and Chemosensory Genes (Dataset S1).

### The Evolution of OR Repertoire Sizes is Governed by Dynamic Evolutionary Changes in Large Subfamilies

Our analysis revealed that although the genomic organization of OR tandem arrays is highly conserved across corbiculate bee lineages, OR evolution is highly dynamic, with multiple taxon-specific OR subfamily expansions. In order to investigate potential localized molecular mechanisms that shape the evolution of the OR gene family, we analyzed the divergence of the 25 OR subfamilies (see above). We found that variation in subfamily size among species was positively correlated with mean subfamily size (Fig. 3a, Pearson coefficient r=0.85, p=0; Orchid bees only: r=0.6, p=0.0013), a common pattern in gene family evolution (Dennis et al. 2017). This result suggests that larger OR subfamilies have a higher gene turnover than smaller gene subfamilies. Accordingly, all subfamilies with one or two genes per species consisted of simple 1:1 orthologous genes across the corbiculate bee genomes and transcriptomes (Fig. S1; Tab. S3).

**Fig. 3.**
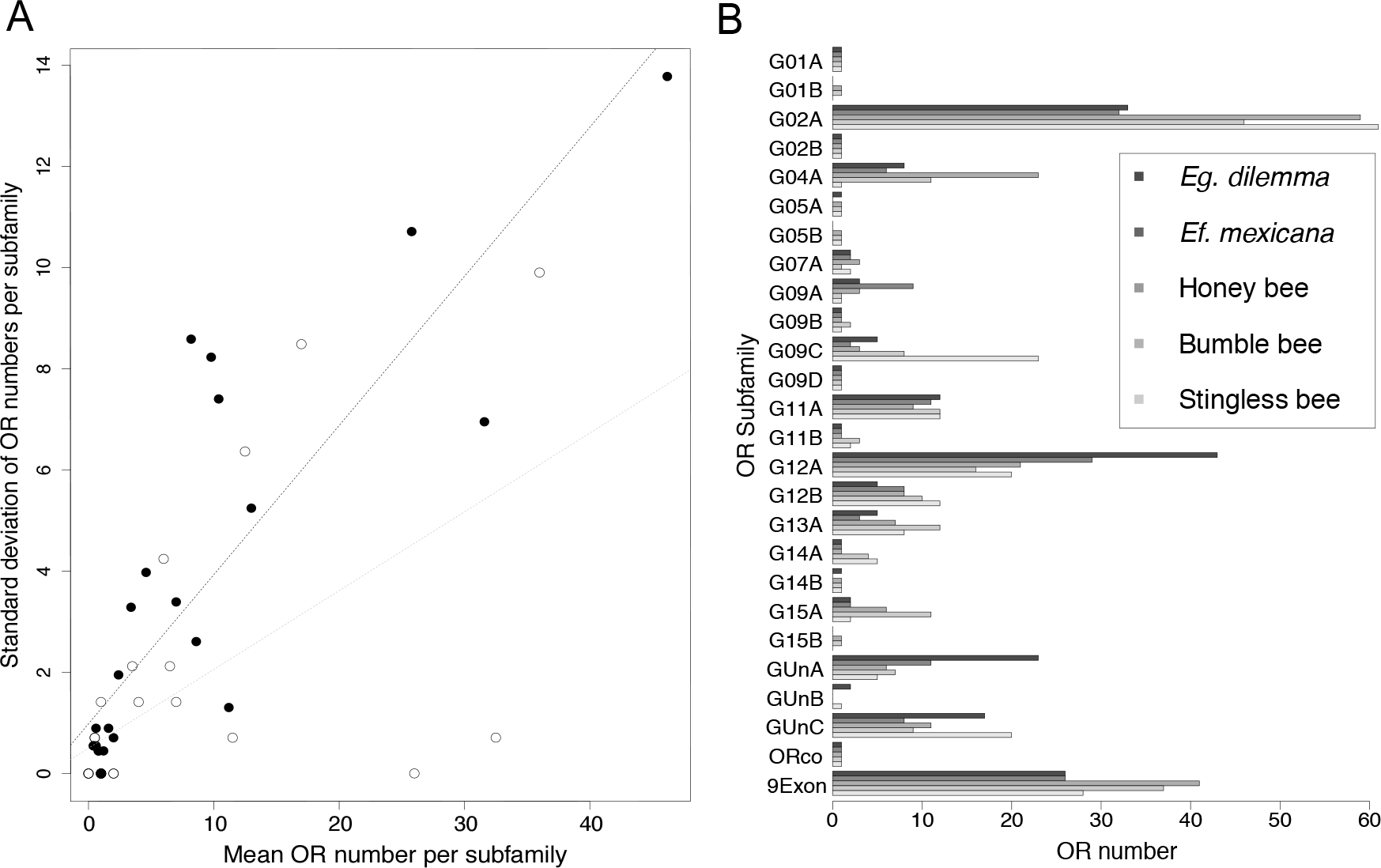
Evolution of OR subfamily size. (**A**) OR subfamily size variation among corbiculates correlates with mean subfamily size. The pattern holds true for all corbiculates (white dots, black dotted line) and the two orchid bee species alone (black dots, grey dotted line). **(B)** OR subfamily repertoire numbers among corbiculate bees. Numbers of ORs in all 25 subfamilies for the five species with whole-genome data are indicated. Pseudogenes are excluded. *Eg. dilemma* and *Ef. mexicana* represent the two orchid bee species. OR subfamily sizes are highly variable between species, while increased subfamily sizes are shared by multiple lineages or occur in single lineages only. 9Exon contains the combined repertoire of subfamilies G04A, G05B, G11A, G11B, and G12A.

Three of the OR subfamilies that exhibited high interspecific variation in OR repertoire size had an expansion in a single bee species. The G04A subfamily was expanded in the honey bee *A. mellifera* (Fig. 2c), the G09A subfamily was expanded in the orchid bee *Ef. mexicana*, and the G09C subfamily was expanded in the stingless bee *M. quadrifasciata* (Fig. 3b, Tab. S3). Several subfamily expansions were observed in two or more species. Although most expansions are not shared by multiple bee species with similar life history traits, the G02A subfamily was expanded in all obligate eusocial bee species we included in our analysis (honey bee, bumble bee and stingless bee; Fig. 3b). Conversely, ORs in the subfamily GUnA and G12A were expanded in orchid bees (Fig. 3b, Tab. S3), which are known for their unique perfume collecting behavior. The expansion we observed in the subfamilies G02A and GUnA correspond to well-supported lineage-specific clusters of paralogs in our phylogenetic analysis (Fig. S1), suggesting that these subfamilies are independent expansions leading to elevated OR numbers. In contrast, more than half of the orchid bee OR expansion in subfamily G12A was shared between the orchid bee species included in our analysis (*i.e*. orthologous copies were present), suggesting that the expansion was already present in their common ancestor.

### Positive Selective Pressures Drive the Evolution of Both Paralogous and Orthologous OR Genes

To investigate the selective pressures that shaped the evolution of the corbiculate bee OR gene family, we directly compared the evolution of ORs with the evolutionary patterns of the remaining major chemosensory gene families, including the GRs, IRs, OBPs, and CSPs. We classified the annotated genome and transcriptome-derived chemosensory genes from all species into orthologous groups consisting of at least four orthologs and/or inparalogs based on our phylogenetic analysis (see Fig. 2b-c for examples of orthologous groups). The resulting 159 orthologous groups (115 OR, 13 GR, 9 IR, 15 OBP, and 7 CSP orthologous groups) contained a total of 1588 chemosensory genes (Dataset S1). We tested each of the orthologous groups for the presence of episodic positive selection using branch-site model based dN/dS estimates (Smith et al. 2015). We identified 54 chemosensory genes (2.1% of all terminal branches) and 57 deeper branches (6.1% of 941 deeper branches) with signatures of positive selection under a 5% false discovery rate (FDR). This corresponds to 4.4% of all branches tested. We found no correlation between divergence time and positive selective pressures (Supplemental Text; Tab. S4). Furthermore, there was no difference in the number of branches under positive selection between the four corbiculate bee tribes (Tab. 1). However, a comparison across gene families revealed a higher proportion of branches under positive selection in the GRs relative to any other gene family in all bee lineages, supporting the hypothesis that GRs exhibit elevated signatures of positive selection across many insect groups (p<0.0001; Fisher’s Exact Test; Supplementary Text) (McBride & Arguello 2007).

**Tab. 1.** Numbers of branches under positive selection for each chemosensory gene family among the four corbiculate bee tribes.

|  |  | Orchid bees | Honey bees | Bumble bees | Stingless bees | Total |
| --- | --- | --- | --- | --- | --- | --- |
| **ORs** | Tested | 1234 | 216 | 194 | 237 | 1881 |
| | $P \leq 0.05$ | 42 (3.4%) | 5 (2.3%) | 3 (1.6%) | 14 (5.9%) | 64 (3.4%) |
| **GRs** | Tested | 116 | 11 | 32 | 15 | 174 |
| | $P \leq 0.05$ | 17 (14.7%) | 2 (18.2%) | 5 (15.6%) | 4 (26.7%) | 28 (16.1%) |
| **IRs** | Tested | 101 | 9 | 9 | 9 | 128 |
| | $P \leq 0.05$ | 5 (5.0%) | 0 (0%) | 0 (0%) | 0 (0%) | 5 (3.9%) |
| **OBPs** | Tested | 178 | 29 | 18 | 12 | 237 |
| | $P \leq 0.05$ | 3 (1.7%) | 4 (13.8%) | 0 (0%) | 0 (0%) | 7 (3.0%) |
| **CSPs** | Tested | 76 | 6 | 5 | 6 | 93 |
| | $P \leq 0.05$ | 1 (1.3%) | 0 (0%) | 0 (0%) | 0 (0%) | 1 (1.1%) |

All branches tested as well as the numbers and percentages of branches under positive selection after 5% FDR correction are represented.

Those branches that exhibited signatures of positive selection were distributed throughout the entire OR gene tree (Fig. S1). Interestingly, only 32% of all branches that we identified to evolve under positive selection belonged to lineage-specific duplications or larger gene expansion events (Tab. S5). Thus, the majority of branches under positive selection correspond to orthologous rather than paralogous gene copies. This pattern held true when we considered only those five species with genomic data. However, when normalized by the total number of the two branch classes (duplication and non-duplication branches, respectively), we found that a higher proportion of all tested duplication branches were under positive selection in comparison to non-duplicated branches (6% vs. 3.9%, Fisher’s Exact Test, p< 0.05, Tab. S5). Together, these results suggest that positive selective pressures play an important role in the evolution and divergence of duplicated OR genes.

In addition, we observed that 10 of the 11 OR subfamilies without any branches under positive selection included simple (single-copy) 1:1 orthologs only (Subfamilies G01A, G01B, G02B, G05A, G09D, G14B, G15A, G15B, GUnB, ORco; Fig. S1). This observation suggests the existence of a core set of OR genes with conserved chemosensory functions across all corbiculate bees.

## Discussion

Our analysis focused on documenting signatures of positive selection as well as changes in the genomic organization of genes in the OR gene family analyzing 1394 ORs spanning multiple evolutionary time scales across the corbiculate bees. Our analysis revealed that the corbiculate bee OR gene family is highly dynamic (Fig. 1, Fig. 2, Fig. S1), a pattern consistent with that observed in other insect lineages. However, the genomic organization we identified in corbiculate bees differs from that previously reported in other insect groups (Guo & Kim 2007; Conceição & Aguadé 2008; Engsontia et al. 2014). First, we found that the genomic organization of OR genes is highly conserved between corbiculate bee lineages, despite sharing a common ancestor 80 million years ago (Tab. S2). Second, OR gene repertoire sizes vary significantly between bee species due to lineage-specific expansions in a few large tandem arrays, some of which correlate with the evolution of obligate eusociality and pheromone communication systems (Fig. 3; Fig. S2). These findings suggest that the highly dynamic evolutionary history of the OR gene family is independent of changes in genomic organization and instead more likely influenced by locally acting molecular mechanisms. Furthermore, we found evidence for elevated rates of positive selection on duplicate ORs (Tab. 1; Tab S6), supporting the main prediction from the birth-death model of gene family evolution. Finally, we identified a fraction of bee ORs under positive selection that have not undergone previous duplication, highlighting the importance of positive selective pressures in OR sequence divergence, possibly in response to ecological adaptation.

### Locally Acting Molecular Mechanisms are the Driving Force of Chemosensory Evolution

Our phylogenetic and genome annotation analysis revealed a high degree of genomic landscape conservation in the OR gene family across the corbiculate bees. This pattern was accompanied by a fast diversification rate of OR genes, indicating that genomic rearrangements had a relatively weak influence relative to the stronger influence of local molecular mechanisms. The high degree of conservation in the overall genomic organization holds true even between corbiculate bee lineages that shared a common ancestor 80 million years ago. This is an unprecedented level of genomic conservatism in OR organization, and is particularly surprising given the diverse factors known to affect genome composition and genome architecture. In fact, the bee lineages we examined exhibit more than two-fold variation in chromosome number (9 in stingless bees to 20 in orchid bees; (Ross et al. 2015)), an order of magnitude of variation in genome size (ranging from 0.25GB in honey bees to 3GB in orchid bees; (Weinstock et al. 2006; Kapheim et al. 2015; Sadd et al. 2015; Brand et al. 2017)), as well as exceptionally high recombination rates (Stolle et al. 2011; Rueppell et al. 2016; Wilfert et al. 2007). Moreover, the high level of conservation stands in sharp contrast to known evolutionary mechanisms that affect genomic organization of OR genes in drosophilid flies, the only other insect group for which the evolution of chemosensory genes has been examined in comparable detail. The genomes of drosophilid flies exhibit complex genome rearrangements leading to a high degree of ortholog OR gene translocation within evolutionary time scales similar to those spanning the bee species we analyzed (Guo & Kim 2007; Nozawa & Nei 2007; Obbard et al. 2012). A similarly dynamic pattern of OR tandem array architecture was previously observed in mammalian genomes (Niimura & Nei 2005). These observations lead to the hypothesis that genomic rearrangements are important in the evolution of chemosensory receptors across vertebrate and invertebrate lineages (Kratz et al. 2002; Sánchez-Gracia et al. 2009; Conceição & Aguadé 2008). In contrast, our results show that the evolution of the OR gene family in corbiculate bees has resulted in highly divergent gene repertoires without undergoing genome rearrangements. The recent annotation of the OR gene family in a larger array of insect genomes suggests that the patterns of OR translocation observed in drosophilid flies are exceptionally high relative to other insect lineages. In fact, non-drosophilid genomes tend to have fewer and larger OR tandem arrays, indicating a less dynamic reorganization of this gene family (Engsontia et al. 2014; Briscoe et al. 2013; Christopher D Smith et al. 2011; Robertson et al. 2010; Engsontia et al. 2008; Smadja et al. 2009). However, singleton ORs are more prevalent in other insect genomes (e.g. Lepidoptera (Engsontia et al. 2014; Briscoe et al. 2013)) relative to corbiculate bees, indicating that OR genomic architecture is less conserved in non-bee lineages. However, additional insect genomes with comparable divergence times are required in order to determine whether the high conservation we observed in corbiculate bee OR genes represents a general or unique pattern.

Our results revealed no correlation between the evolution of genomic organization and genome size. In fact, genome size of most corbiculate bees resembles those reported for drosophilid flies (Bosco et al. 2007; Weinstock et al. 2006). Instead, recent comparisons of the genomes of honey bees, bumble bees, and orchid bees revealed high conservation of genome synteny between these species (Stolle et al. 2011; Brand et al. 2017), indicating that conserved genomic architecture of OR tandem arrays may result from a low precedence of genome rearrangements in corbiculate bees. Indeed, our analysis revealed that synteny among OR genes in orchid bees is almost perfectly conserved even within highly dynamic genomic areas that are dominated by gene duplication (Supplementary Text). Despite the high level of conservation in the genomic organization reported here, we find a high degree of OR gene divergence and turnover among corbiculate bees. This result suggests that chemosensory gene evolution in corbiculate bees, and perhaps insects in general, is shaped by local rather than global mechanisms of genome evolution.

### Large Tandem Arrays are Hotspots for the Evolution of Novel ORs

The high rate of variation we observed in OR repertoire size among lineages, along with the prevalence of lineage-specific gene expansions in large OR tandem arrays, suggests that large gene subfamilies play an important role in the evolution of novel OR genes in corbiculate bees. Insect OR repertoires are highly dynamic between lineages in comparison to other chemosensory gene families (Sánchez-Gracia et al. 2009; Croset et al. 2010). Our results confirm this pattern for corbiculate bees. However, we find that these expansions are driven by lineage-specific expansions within large tandem arrays harboring entire OR subfamilies. This result is congruent with a recent phylogenetic analysis that showed that gene duplication and pseudogenization events of OR genes in Hymenoptera are often found in large subfamilies (Zhou et al. 2015). The increased gene turnover rate could be influenced by elevated rates of genome replication errors resulting from non-allelic homology of a large number of closely related paralogous loci in close proximity within large tandem arrays (Hastings et al. 2009; Perry et al. 2008; 2006). Nevertheless, other factors such as transposable element activity and the timing of DNA replication can influence segmental duplication rates (Hastings et al. 2009; Cardoso-Moreira et al. 2011). Independent of the underlying genomic mechanism, our results suggest that large OR tandem arrays are a major source of novelty in chemosensory receptors in corbiculate bees and potentially other Hymenoptera. Accordingly, large tandem arrays are likely an important source of functional novelty associated with the detection of novel molecules or blends of chemical compounds. Hence, it is possible that the gene expansions we identified coincide with key evolutionary innovations such as complex social chemical communication in the obligate eusocial honey bees, bumble bees and stingless bees or perfume-collecting behavior in orchid bees.

It has been hypothesized that OR gene family size is correlated with the degree of complexity in the chemical communication system displayed by eusocial Hymenoptera (LeBoeuf et al. 2013). Although our data do not support this general trend, we find a positive correlation between OR gene subfamily size and the presence of obligate eusociality. For instance, the OR subfamily G02A is expanded in all three obligate eusocial bee species analyzed, relatively to the smaller size exhibited by the solitary to weakly social orchid bees. Moreover, the OR subfamily G02A is expanded in ants but not in the facultative social halictid bee *Lasioglossum albipes*, which exhibits subfamily G02A gene numbers comparable to orchid bees (Subfamily L in (Engsontia et al. 2015; Zhou et al. 2015)). Thus, it is possible that ORs in subfamily G02A play a central role in the olfactory detection and processing of pheromones involved in social behavior. Similarly, an expansion of the ‘9-exon’ subfamily (here subfamilies G04A, G05B, G11A, G11B, and G12A; Fig. 2a; Fig. 3b; Fig S1) was hypothesized to be important in the evolution of eusociality due to a role of ‘9-exon’ ORs in the detection of social pheromones in ants (namely cuticular hydrocarbons, (Christopher D Smith et al. 2011; Chris R Smith et al. 2011; McKenzie et al. 2016; Slone et al. 2017)). We find an expansion of the ‘9-exon’ subfamily in the eusocial honey bees and bumble bees but not eusocial stingless bees. This observation contrasts with the previous suggestion that the expansion of this OR subfamily was concomitant with the evolution of eusociality in Hymenoptera.

However, it is possible that ORs involved in social pheromone communication evolved independently in different Hymenoptera lineages. A recently published study could show that cuticular hydrocarbons involved in social communication in ants are ligands to ORs in multiple subfamilies (Slone et al. 2017). Future studies should investigate the possibility that olfactory adaptations to social behavior evolved through multiple analogous routes in social Hymenoptera (Woodard et al. 2011; Kapheim et al. 2015).

In addition, we found two OR subfamilies that exhibit an expansion in the orchid bee lineage (namely GUnA and G12A). While large expansions were in general species-specific clusters of paralogous ORs (*e.g*. Fig. 2c; Fig. S1), more than half of the ORs in the orchid bee expansion of subfamily G12A were orthologous ORs found in all the seven species included in our analysis. This shared expansion may correspond to an orchid bee specific olfactory specialization. Since the only known olfactory behavior clearly distinguishing orchid bees from all other corbiculate bees is male perfume-collecting behavior (Roubik & Hanson 2004; Michener 2007), it is possible that the G12A subfamily is involved in the detection and encoding of these exogenous compounds. Functional analyses are required to investigate the olfactory coding mediated by members of this OR subfamily.

### Selective Patterns of Corbiculate Bee Chemosensory Genes Support the Birth-Death Model of Gene Family Evolution

It has been proposed that chemosensory gene families evolve under a birth-death process where duplications of existing genes give rise to new gene copies (gene ‘birth’) whereas pseudogenization purges genes from the genome (gene ‘death’; (Nei & Rooney 2005)). The birth-death model predicts a relaxation of purifying selective pressures after duplication allowing subsequent diversification of one or both copies through positive selection for newly accumulated beneficial mutations. Accordingly, the evolution of highly dynamic chemosensory gene families is likely driven by the action of positive selection. Nevertheless, while previous studies detected a high impact of gene duplication and loss on chemosensory gene family evolution, evidence for diversification through positive selection is limited but has been reported for paralogous chemosensory receptors in several insect species (Guo & Kim 2007; Tunstall et al. 2007; Gardiner et al. 2008; McBride & Arguello 2007; Engsontia et al. 2014; Zhou et al. 2015; Engsontia et al. 2015; Smadja et al. 2009; Croset et al. 2010).

Our analysis provides evidence congruent with the selective pressures predicted by the birth-death model of chemosensory gene family evolution. We show that positive selective pressures are stronger on duplicated chemosensory genes than in orthologous chemosensory genes across corbiculate bees. It has been proposed that non-homologous gene conversion can have a moderate negative impact on the inference of selection when using maximum likelihood methods, particularly due to an increased false-positive error rate between recent duplicates (Casola & Hahn 2009). However, previous analyses have shown that non-homologous gene conversion is rare in insect chemosensory genes, the potential effects on gene tree inference and selection analysis tends to be small (Almeida et al. 2014) Moreover, the unlikely presence of false positives due to gene conversion would further highlight the importance of positive selection on orthologous chemosensory genes in corbiculate bees. The observed higher proportion of branches under positive selection is in contrast to previous analyses of insect chemosensory genes and might be partly explained by the former use of less powerful algorithms, reliance on a priori expectations, incomplete testing, or differences in taxon sampling (Martin D Smith et al. 2015). It is thus unclear whether the disparity between studies reflects technical differences or differences in chemosensory gene evolution between different groups of insects. Regardless, our findings suggest a non-negligible impact of positive selective pressures on chemosensory gene evolution in corbiculate bees.

### Diversification of Orthologous ORs through Positive Selection Drives Divergence in Chemosensory Systems

Our selection analysis indicates that positive selection is involved in the divergence of orthologous ORs, providing an evolutionary mechanism for the diversification of chemosensory genes independent of gene duplication. We found 54 ortholog chemosensory genes and deeper branches that exhibit signatures of positive selection. Moreover, our analysis using three pairs of sympatric species revealed the presence of orthologous chemosensory genes evolving under positive selection irrespective of divergence times (0.15 - 13 my). This result suggests that positive selection is an important evolutionary force driving the diversification of orthologous chemosensory genes in corbiculate bees. Previous studies have highlighted the importance of divergence in orthologous chemosensory receptor genes in the divergence of insect species and/or populations (Smadja et al. 2012; Eyres et al. 2017; Leary et al. 2012). For example, Leary et al. (2012) showed that positive selection of specific amino acid substitutions in OR orthologs led to novel receptor functions through changes in ligand binding affinities in *Ostrinia* moths. Under this model, orthologous OR genes undergo functional divergence driven by positive selection in the absence of a duplication event. While OR gene duplications are more common in large subfamilies in corbiculate bees, this mechanism might be important for the evolution of interspecific functional divergence in smaller subfamilies. Thus, our results suggest that positive selection is not only important in driving divergence between OR duplicates within taxa, but also orthologous ORs between taxa. It is also possible that a combination of these two mechanisms drives the evolution of chemosensory genes in other insect lineages.

### Evidence for ORs with Core Chemosensory Function in Corbiculate Bees

We detected the existence of multiple highly conserved OR lineages that appear to be subject to strong purifying selection. This observation supports the existence of a core set of OR genes with conserved functions across all corbiculate bees and perhaps other more distantly related bee lineages. We detected 10 OR subfamilies and 14 orthologous groups within larger subfamilies that solely consisted of 1:1 orthologs. While most OR subfamilies and orthologous groups were characterized by taxon-specific duplications and divergence driven by positive selective pressures, our results suggest that these OR lineages are highly conserved throughout at least 80 million years of evolution. This observation implies that corbiculate bee genomes contain a core set of olfactory genes with conserved chemosensory functions. In fact, high conservation of OR genes is typical for some species with highly similar natural history (Neafsey et al. 2015; McBride & Arguello 2007; Nozawa & Nei 2007). Detailed functional analysis of these core OR genes may elucidate the role of conserved olfactory functions in corbiculate bees of ecological importance.

## Conclusion

Odorant receptors (ORs) have been hypothesized to play an important role in chemosensory adaptation, for example by enabling the evolution of novel pheromone detection systems associated with social communication or mate attraction (Hansson & Stensmyr 2011; Smadja & Butlin 2009). However, little is known about the molecular mechanisms driving the evolution of insect OR gene families. Our analysis of the OR gene family in an important lineage of insect pollinators, the corbiculate bees, elucidated the role of positive selective pressures and genomic organization on the evolution of the OR gene family.

We show that OR gene family size is highly dynamic between corbiculate bee species. In contrast, the underlying genomic organization of OR genes is highly conserved among lineages over at least 80 million years. Consistent with these findings, ORs are arranged in tandem arrays located in close proximity along the genome. Moreover, the genomic location correlates with the phylogenetic relationships among genes that fall in 25 well-supported OR subfamilies. We find that a few large subfamilies contain a considerable amount of lineage-specific gene duplications, explaining most of the variation in gene family size. Further, we show that ORs evolve under strong positive selective pressures, with a higher impact on duplicated genes. Our study suggests that locally acting molecular mechanisms are the driving force of OR gene family evolution in corbiculate bees. Moreover, we provide empirical support for the longstanding hypothesis that positive selection is an important mechanism in the evolution of duplicated OR genes in insects. These results elucidate the OR gene family evolution in corbiculate bees and extend the current understanding of OR gene family evolution to other insect groups.

## Methods

### Sampling and Sequencing

Sampling and sequencing procedures followed Brand et al. 2015. Briefly, males of the four orchid bee species *Euglossa flammea*, *Eg. imperialis*, *Eulaema bombiformis* and *El. meriana* were sampled at the La Gamba Field Station in Costa Rica using different chemical baits (Pokorny et al. 2013). Antennal transcriptomes for each species were based on pools of the antennae of 20 individuals for the *Euglossa* species and 5 individuals for the *Eulaema* species. Bees were chilled on ice and the antennae of each torpid male were dissected using sterile forceps and transported either on liquid nitrogen or RNA-later (Thermo Fisher Scientific, Waltham, MA) and kept frozen until RNA-extraction.

Total-RNA was extracted and quantified as described (Brand et al. 2015). Barcoded cDNA libraries were prepared using the NEBnext Ultra RNA Library Prep kit for Illumina (New England Biolabs, Ipswich, MA) following the manufacturers protocol and sequenced in two pools of two species on one HiSeq 2500 (*Euglossa*) and one HiSeq 4000 (*Eulaema*) lane for 2x100bp cycles. (Raw sequence reads are available at the NCBI Sequence Read Archive under BioSample access number PRJNA387619).

### *De novo* transcriptome assembly and transcript recovery

Identical raw reads were merged using fastuniq v1.1 (Xu et al. 2012) and subsequently trimmed by trim_galore v0.3.7 (Babraham Bioinformatics) if sequencing primers or low-quality bases (Phred-score <20) were detected. The pre-processed reads of each species were used for *de novo* transcriptome assembly following the meta-assembly approach developed in Brand et al. 2015 using the Trinity (Grabherr et al. 2011; Haas et al. 2013) and SOAPdenovo-trans v2 (Xie et al. 2014) assemblers. Briefly, each of the two assemblers was run with multiple parameter sets controlling assembly stringency resulting in 25 assemblies per species. Trinity settings were set as described (Brand et al. 2015), whereas SOAPdenovo-trans parameters −e and −d were set to 0, 2, 3, or 5 and 1, 3, 5, or 7, respectively and run in all possible combinations. While Trinity was run with the default single-k-mer mode, SOAPdenovo-trans was run in multi-k-mer mode with nine different k-values (25, 31, 37, 43, 49, 59, 69, 79 and 89).

### Annotation of chemosensory genes

Chemosensory genes of five gene families (ORs, IRs, GRs, OBPs, and CSPs) were annotated in the newly sequenced antennal transcriptomes of the orchid bee species *Eg. flammea*, *Eg. imperialis*, *El. bombiformis*, and *El. meriana*, as well as the publicly available genomes of the orchid bee species *Eg. dilemma* (Brand et al. 2017) and *Eufriesea mexicana* (Kapheim et al. 2015), and the stingless bee *Melipona quadrifasciata* (Kapheim et al. 2015). Additionally, the CSP gene family was annotated for the bumble bee *Bombus terrestris* (Sadd et al. 2015). All four *de novo* transcriptomes were annotated using a homology-based iterative tBLASTn approach as previously described (Brand et al. 2015). In order to annotate the genomes, tBLASTn was used to detect scaffolds harboring chemosensory genes with an e-value cutoff of 10e-6 based on gene family specific query libraries comprised of published bee, wasp, ant and dipteran OR, IR, GR, CSP and OBP sequences (Robertson & Wanner 2006; Brand et al. 2015; Sadd et al. 2015; Forêt & Maleszka 2006; Croset et al. 2010; Vieira & Rozas 2011; Zhou et al. 2012). Subsequently, the queries were used to predict exon-intron boundaries with exonerate (Slater & Birney 2005) in a scaffold specific manner. All annotations were curated manually and corrected if needed. All annotations can be found in Dataset S1.

### Phylogenetic reconstruction of chemosensory gene families

For each chemosensory gene family we estimated a maximum likelihood (ML) gene-tree using RaxML (Stamatakis et al. 2005) to infer the genealogical histories of the candidate corbiculate gene family members. On that account, gene family specific protein sequence alignments were constructed using MAFFT (Katoh et al. 2002; Katoh & Standley 2013) applying the L-INS-I algorithm with the–maxiterate option set to 1000 (Katoh et al. 2005). Based on the resulting alignments ML trees were inferred under a JTT + G substitution model (Brand et al. 2015) for each family with 20 independent ML searches and 100 bootstrap replicates.

### Analysis of the genomic organization of chemosensory gene families

Based on genomic annotations of *Eg. dilemma*, *Ef. mexicana*, *M. quadrifasciata* (this study), *A. mellifera* (Robertson & Wanner 2006), and *B. terrestris* (Sadd et al. 2015), we analyzed the evolutionary history of genomic structure of chemosensory gene families within the corbiculate bees. Therefore, we applied a reciprocal blast approach that compares the genomic location of all chemosensory gene family tandem arrays and singletons to the genomes of all other species. We required both 200kb flanking regions to be the reciprocal best blast hit (Rivera et al. 1998) in order to call homology of the genomic regions. In case annotations were located less than 200kb away from the end of a scaffold, we used the maximum flanking region length possible, which never corresponded to less than 100kb. The location of ORs in the resulting genomic regions in combination with the gene family phylogeny was used to classify all ORs into subfamilies. Subfamilies were named after the linkage group they were located on in the honey bee genome. Letters were used to allow for the unique indication of subfamilies located on the same linkage group (*e.g*. G12B is the second subfamily found to be located on linkage group 12 of the honey bee). In case a gene family was located on a honey bee scaffold that could not be assigned to a linkage group, it was named ‘Un’, in short for ‘unassigned’.

In a second approach, we used the OR gene family phylogeny to infer the evolutionary histories of micro-genomic synteny within tandem arrays. Orthologs and paralogs were directly inferred from the ML phylogenies and subsequently mapped to respective genomic coordinates. Since more distantly related species share less orthologous than paralogous gene copies (*e.g*. (Robertson & Wanner 2006; Zhou et al. 2015)), we focused on the two orchid bees *Eg. dilemma* and *Ef. mexicana* for this analysis.

### Selection analysis

Based on the phylogenetic reconstructions including all genome and transcriptome derived genes (see above), we defined monophyletic orthologous groups for each gene family consisting of four or more ortholog copies including in-paralogs that were used for all subsequent selection analyses. In order to identify genes under positive selection, we applied the adaptive branch-site relative effects-likelihood (aBSREL) algorithm as implemented in HYPHY (Pond et al. 2005) to each subgroup to identify branches with dN/dS ratios significantly larger than 1, indicating positive selection along these branches (Martin D Smith et al. 2015). P-values for each aBSREL run were corrected for multiple testing using a false-discovery-rate (FDR) of 5%. In order to analyze patterns of recent gene family divergence in the orchid bees, we conducted pairwise dN/dS analyses for the *Eulaema* species pair and the two *Euglossa* species pairs. Therefore, we extracted orthologous copies from the gene family phylogenies and performed pairwise nucleotide sequence alignments for each orthologous pair. Each alignment was subsequently used for ML estimation of the pairwise dN/dS ratio (model M1) in codeml (Bielawski & Yang 2004; Yang 2007). dN/dS ratios for each pairwise comparison were tested for significant deviations from the neutral expectation (dN/dS=1; model M0) using likelihood-ratio tests.

## Data Access

All gene annotations are included as Supplemental Material. Raw sequence reads are available at the NCBI Sequence Read Archive under BioSample access number PRJNA387619.

## Acknowledgements

We thank Michelle Stitzer, Judy Wexler, Mascha Koenen, the Ramírez lab, and two anonymous reviewers for valuable comments on the manuscript. We thank Hugh Robertson for sharing chemosensory gene sequences of the honey bee and bumble bee. P.B. was supported by a fellowship of the German academic exchange service (Deutscher Akademischer Austauschdienst, DAAD) for parts of the project. SRR received support from the David & Lucile Packard Foundation and the National Science Foundation (DEB-1457753). This work used the Vincent J. Coates Genomics Sequencing Laboratory at UC Berkeley, supported by NIH S10 Instrumentation Grants S10RR029668, S10RR027303, and OD018174.

## Disclosure Declaration

The authors report no conflicts of interest.

